# Rapid generation of single-insertion transgenics by Tol2 transposition in zebrafish

**DOI:** 10.1101/2023.10.11.561928

**Authors:** Miglė Kalvaitytė, Sofija Gabrilavičiūtė, Darius Balciunas

## Abstract

The Tol2 transposable element is the most widely used transgenesis tool in zebrafish. However, its high activity almost always leads to multiple unlinked integrations of the transgenic cassette in F_1_ fish. Each of these transgenes is susceptible to position effects from surrounding regulatory landscape, which can lead to altered expression and, consequently, activity. Scientists therefore must strike a balance between the need to maximize reproducibility by establishing single-insertion transgenic lines and the need to complete experiments within a reasonable timeframe. In this article, we introduce a simple competitive dilution strategy for rapid generation of single-insertion transgenics. By using *cry:BFP* reporter plasmid as a competitor, we achieved a nearly fourfold reduction in the number of the transgene of interest *(TgOI*) integrations while simultaneously increasing the proportion of single-insertion F_1_ generation transgenics to over 50%. We also observed variations in *TgOI* expression among independent single-insertion transgenics, highlighting that the commonly used ubiquitous *ubb* promoter is susceptible to position effects. Wide application of our competitive dilution strategy will save time, reduce animal usage, and improve reproducibility of zebrafish research.

**Summary:** Competitive dilution of Tol2 transgenesis vectors facilitates isolation of single-insertion transgenic zebrafish lines in the F_1_ generation, reducing animal usage, improving reproducibility, and saving time.

## INTRODUCTION

Transposon-based systems enable efficient and precise integration of foreign DNA into the zebrafish genome (Balciunas et al., 2006; Davidson et al., 2003; Emelyanov et al., 2006; Kawakami and Noda, 2004; Kawakami et al., 2004b; Quach et al., 2015; Sato et al., 2007). Compared to conventional DNA microinjection methods (Lawson and Weinstein, 2002; Thermes et al., 2002), transposons offer two major advantages: a high germline transmission rate and single-copy (non-concatemerized) integration of the transgene. Originally isolated from the medaka fish genome, the Tol2 transposon was recognized for its high activity and integration efficiency, leading to its rapid adoption as the primary transgenesis tool for zebrafish (Balciunas et al., 2006; Kawakami et al., 2004a; Koga et al., 1996). While other tools for zebrafish transgenesis, such as I-SceI (Thermes et al., 2002), PhiC31 (Groth et al., 2004; Roberts et al., 2014), Sleeping Beauty (Davidson et al., 2003) and Ac/Ds (Chong-Morrison et al., 2023; Emelyanov et al., 2006; Quach et al., 2015) are available, Tol2 remains the leading choice due to its remarkably robust and high germline transmission rate (Balciunas et al., 2006; Kawakami et al., 2004a).

The widespread application and growing demand for streamlined genetic modifications in zebrafish have driven significant advancements in Tol2 transgenesis. *Definition of minimal* length of inverted terminal repeat (*ITR*) sequences required for Tol2 transposition (*miniTol2*) effectively reduced vector size by 3 kb (Balciunas et al., 2006; Urasaki et al., 2006). Development of the Tol2kit further facilitated broad adoption of Tol2 by introducing a collection of Gateway-compatible vectors containing commonly used elements such as promoters and reporters (Kwan et al., 2007). This library of compatible components has been recently thoroughly expanded (Kemmler et al., 2023). In addition, many laboratories possess their own libraries of constructs, making it very likely that Tol2 will remain the transgenesis tool of choice for zebrafish.

The Tol2 transposon usually leads to the precise integration of a single-copy of the transgene at a random location. However, it is common for F_1_ fish to have up to 10 or more transposon integrations spread throughout the genome (Balciunas et al., 2006; Distel et al., 2009; Kawakami and Noda, 2004; Kawakami et al., 2004a; Nagayoshi et al., 2008; Zhang et al., 2019). This feature has been extensively utilized in gene- and enhancer-trap approaches, where multiple integrations of the trap cassette, combined with the high rate of germline transmission, enable selection for rare “trapping” events (Balciuniene et al., 2013; Chan et al., 2019; Clark et al., 2011a; Davison et al., 2007; Grajevskaja et al., 2018; Kawakami et al., 2010; Nagayoshi et al., 2008; Parinov et al., 2004; Rehn et al., 2011). High transposon load, however, becomes a drawback when aiming to establish transgenic lines with specific and reproducible expression of the transgene. Given that each transgene is likely to be susceptible to position effects (Grajevskaja et al., 2013; Jaenisch et al., 1981), different integrations almost inevitably display differences in transgene expression strength and pattern (Hans et al., 2011; Roberts et al., 2014). With a standard generation time of about three months, isolation of single-insertion zebrafish lines by outcrossing multiple-integration F_1_s can easily take a year. Balancing the urgency of experimental timelines with a time-consuming nature of purifying multicopy lines presents a challenge, potentially leading to issues of reproducibility due to variation in transgene copy number and composition in subsequent generations.

To expedite the generation of single-insertion zebrafish transgenic lines, we modified the standard Tol2-mediated transgenesis protocol by diluting the vector carrying the transgene of interest (*TgOI*) with a “competitor” Tol2 vector marked with a fluorescent reporter for straightforward (counter) selection. This competition led to a reduction in the number of integrated *TgOI* cassettes into the zebrafish genome, allowing us to successfully generate multiple independent single-insertion transgenic lines in the F_1_ generation, bypassing the need for multi-generational breeding.

## RESULTS AND DISCUSSION

### Competitive dilution strategy for single-insertion transgenics

In standard Tol2-mediated transgenesis protocol, a mixture of 20-30 pg of plasmid carrying the *TgOI* and 25 pg of Tol2 mRNA is injected into single-cell stage zebrafish embryos. This results in up to 70% of injected F_0_ fish becoming germline transmitting founder fish passing on *TgOI* insertions to their progeny (Balciunas et al., 2006; Balciuniene et al., 2013; Clark et al., 2011b; Kawakami et al., 2004a). We hypothesized that by diluting the target vector with another one, while maintaining the same mass ratio of Tol2 donor vector and Tol2 mRNA, we would create a competition between the two vectors for the Tol2 transposase. This should lead to a reduced number of *TgOI* insertions transmitted to the F_1_ generation while the germline transmission rate of both transgenes in total should not be affected. To test this hypothesis, we diluted the *TgOI* expression vector with a simple *cry:BFP* reporter vector in a 1:4 mass ratio and injected this mix into the single-cell-stage embryos along with Tol2 mRNA (Fig. 1A). Germline transmitting founder fish were selected through incross (Fig. 1B). *TgOI* copy number in the F_1_ generation was assessed by analysing the ratio of TgOI-expressing F_2_ embryos (Fig. 1C) and/or by quantitative PCR (qPCR).

**Fig. 1.**
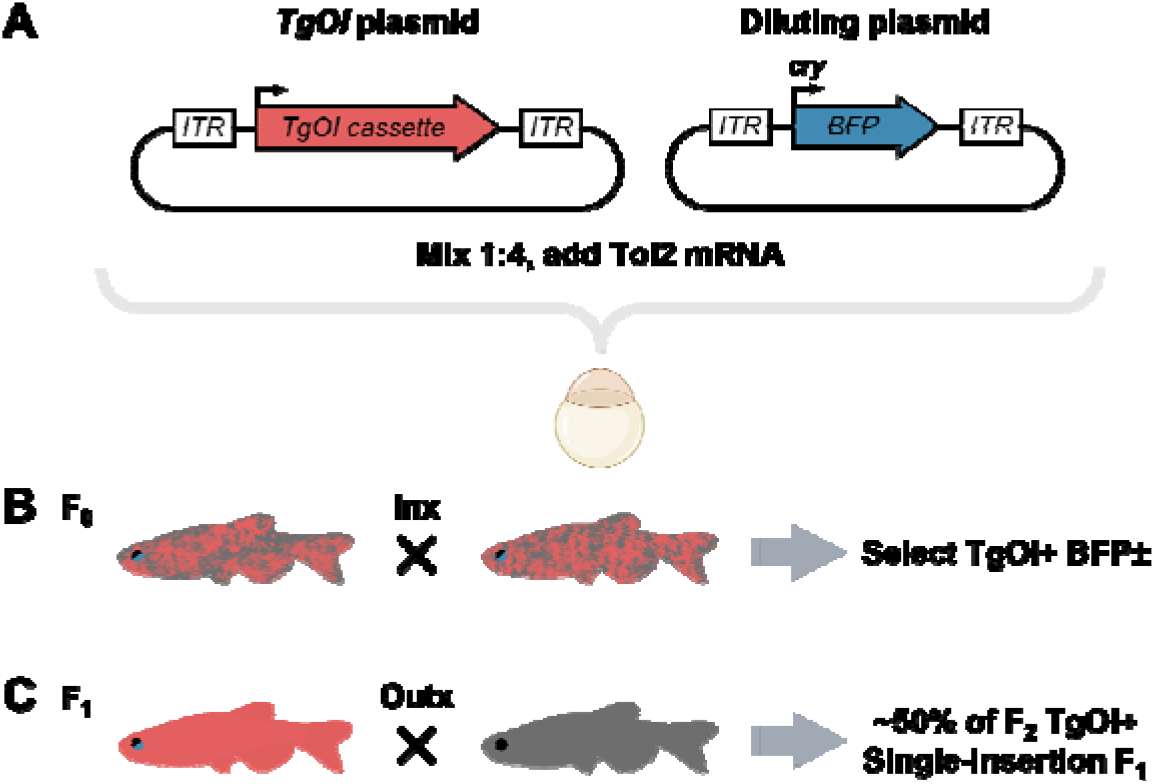
Schematic representation of the competitive dilution strategy to generate single-insertion transgenics in F_1_ generation. (A) The *TgOI* plasmid is mixed with diluting plasmid in a 1:4 mass ratio and co-injected into single-cell stage embryos together with Tol2 transposase mRNA. (B) The F_0_ adult fish are incrossed and screened for germline transmission. (C) TgOI-positive F_1_ generation fish are outcrossed to wild type fish and the offspring are analysed for the expression of *TgOI*. If ∼50% of the offspring carry the *TgOI*, the analysed F_1_ is considered to be a single-insertion transgenic.

### Germline transmission and germline mosaicism of the diluted transgenes

To test our hypothesis, we constructed *miniTol2* expression vectors (Balciunas et al., 2006) containing a *loxP-mRFP-STOP-loxP* cassette driven by the *ubiquitin (ubb)* promoter (Mosimann et al., 2011). Downstream of these cassettes, we inserted two different transgenes: *nup35-eGFP-BLRP* (Amin et al., 2014) to create plasmid pDB986 (Fig. 2A), and *tcf21* coding sequence fused with a triple HA tag at 3’ end to create plasmid pMK17 (Fig. 2B). For dilution, we used a *miniTol2* vector with the *gamma-crystallin* promoter driving the BFP reporter expression (plasmid pDB815) (Fig. 2C). The *TgOI* plasmids were individually mixed with the diluting plasmid in 1:4 mass ratio. The resulting plasmid mixtures (25 pg/embryo) were co-injected with Tol2 mRNA (25 pg/embryo) into the yolks of single-cell-stage embryos following established protocol (Balciunas et al., 2006; Balciuniene and Balciunas, 2013). As a control experiment, we also injected a mixture of 25 pg of undiluted pMK17 with 25 pg of Tol2 mRNA into the yolks of single-cell-stage embryos. Over 90% of embryos from each injection displayed mosaic expression of mRFP and were raised to adulthood for further analysis.

**Fig. 2.**
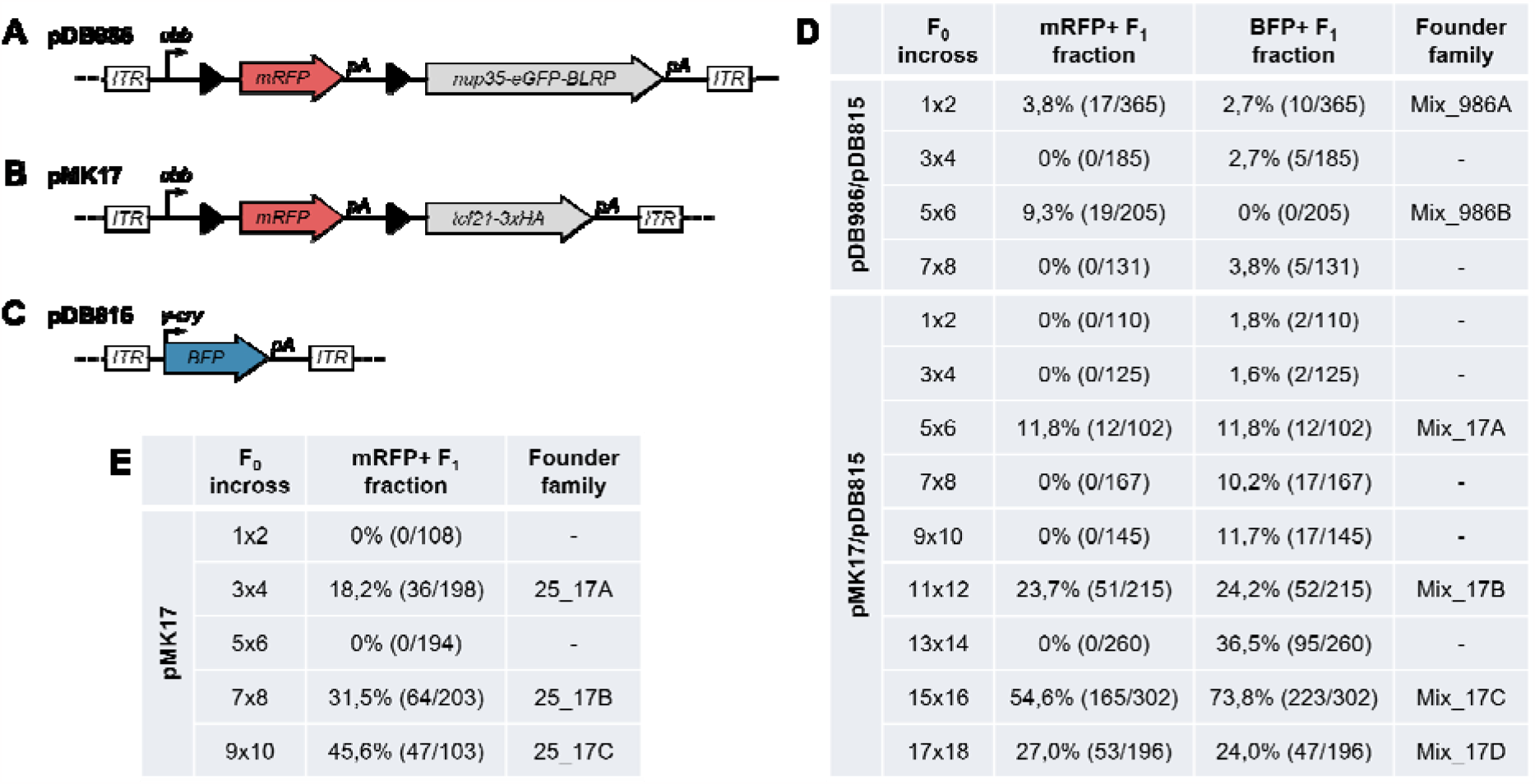
Germline transmission of competitively diluted transgenes. (A-C) Schematic representation of the constructs used in this study: (A) pDB986 (plasmid size: 10000 bp; transgene size: 7060 bp); (B) pMK17 (plasmid size: 8822 bp; transgene size: 5882 bp); (C) pDB815 (plasmid size: 4656 bp; transgene size: 1716 bp). *ITR* – *miniTol2* inverted terminal repeat; pA – polyA. Solid triangles indicate loxP sites. (D) F_0_ fish injected with either pDB986/pDB815 or pMK17/pDB815 plasmid mix were incrossed and their offspring were scored for mRFP and BFP fluorescence. Pair of F_0_ fish transmitting *TgOI* were named as distinct founder family. (E) F_0_ fish injected with 25 pg of pMK17 plasmid were incrossed and their offspring were scored for mRFP fluorescence. Pair of F_0_ fish transmitting TgOI were named as distinct founder family.

To test for germline transmission, we incrossed F_0_ fish from each injection and scored F_1_ embryos for BFP and mRFP fluorescence. Among the F_0_ fish injected with the pDB986/pDB815 plasmid mix (Mix_986 injection), 2 out of 4 incrosses produced mRFP-positive progeny and 3 out of 4 incrosses produced BFP-positive progeny. Similarly, for the pMK17/pDB815 plasmid mix (Mix_17 injection), 4 out of 9 incrosses produced mRFP-positive progeny while all nine incrosses produced BFP-positive progeny (Fig. 2D). In comparison, among the F_0_ fish injected with 25 pg of the pMK17 plasmid (25_17 injection), 3 out of 5 incrosses transmitted the *TgOI* to their offspring (Fig. 2E). While we observed slight variation in germline mosaicism rates among different injections, effectiveness of transgenesis remained consistent with our previous observations (Balciunas et al., 2006; Balciuniene et al., 2013).

### Assessment of *TgOI* copy number in F_1_ generation

To determine the approximate copy number of the transgene, two random mRFP-positive F_1_ fish from Mix_986A and Mix_986B founder families were outcrossed to wild-type fish, and their offspring were visually screened. If approximately 50% of the F_2_ embryos exhibit TgOI-positive traits, then it is likely that the F_1_ parent has a single transgenic insertion. Results showed that the F_1_ fish from Mix_986A founder family transmitted *TgOI* to 93 out of 177 embryos (52,5%), while the F_1_ fish from Mix_986B founder family transmitted *TgOI* to 9 out of 14 embryos (64,3%). These two independent F_1_ fish from different founder families were used to establish stable transgenic zebrafish lines *tpl133Tg* and *tpl134Tg* (Fig. S1), both of which have shown constant single-insertion Mendelian inheritance for multiple generations.

For Mix_17 experiment we pursued a more comprehensive quantitative analysis of *TgOI* copy number variation within the F_1_ generation. The offspring of F_0_ fish, injected with a pMK17/pDB815 plasmid mix, were visually screened for mRFP and BFP fluorescence. 11 to 20 individual mRFP-positive F_1_ embryos from each founder family were imaged at 3 dpf using a fluorescence microscope and collected at 5 dpf for further analysis. For copy number determination using qPCR, we selected 5 embryos from each incross, spanning the full range of mRFP expression levels to capture variation in the *TgOI* copy number.

qPCR analysis revealed that a large fraction of analysed F_1_s from three different Mix_17 founder families (Mix_17A, Mix_17C and Mix_17D) possessed single-insertion of *TgOI*, while the fourth founder family (Mix_17B) did not produce any single-insertion embryos (Fig. 3A). By analysing mRFP expression level of imaged mRFP-positive F_1_ embryos, we determined that within the Mix_17 founder families, 3 out of 4 transmitted a single *TgOI* copy to more than 50% of their offspring (Fig. S2). Meanwhile, copy number of the *BFP* transgene was 2-4 times higher than that of the *TgOI*. Importantly, some of the F_1_s were negative for lens BFP expression, indicating absence of the diluting transgene (Fig. 3A).

**Fig. 3.**
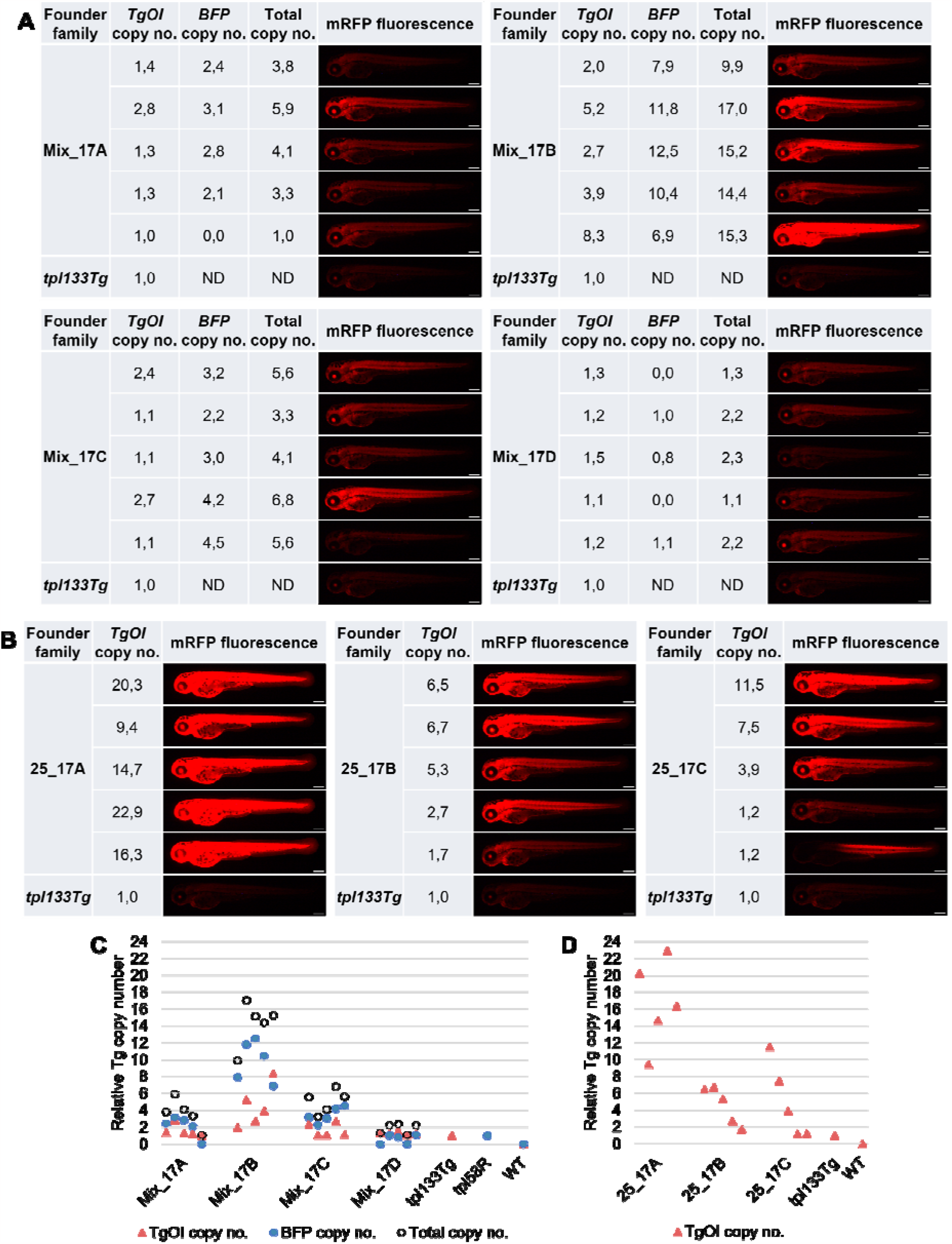
Copy number and strength of transgene expression in F_1_ embryos. (A) Estimated *TgOI* and BFP transgene copy number and mRFP expression in F_1_ embryos from 4 different founder families injected with a mix of pMK17/pDB815 plasmids. (B) Estimated *TgOI* copy number and mRFP expression in F_1_ embryos from 3 different founder families injected with 25 pg of pMK17 plasmid. (C) *TgOI* and BFP transgene copy number variation in F_1_ embryos from different founder families injected with pMK17/pDB815 plasmids mix. Total copy number was determined by summing the copy numbers of both the *TgOI* and BFP transgenes. (D) *TgOI* copy number variation in F_1_ embryos from different founder families injected with 25 pg of pMK17 plasmid. A heterozygous tpl133Tg embryo was used as single copy control for *TgOI*, and heterozygous tb ×5a^*tpl58R*^ gene trap line, containing the gamma-crystalline:BFP expression cassette (Grajevskaja et al., 2018), was used as single copy control for BFP transgene. Scale bars in A,B: 250 μm.

Compared to transgenics obtained from Mix_986 and Mix_17 injections, the majority of F_1_ embryos arising from 25_17 injection displayed much brighter mRFP expression, likely indicating higher *TgOI* copy number (Fig. S3). qPCR analysis on three different founder families revealed that the offspring of 25_17A founder family inherited more than 9 copies of the *TgOI*, while the 25_17B and 25_17C founder families, on average, transmitted ∼5 copies of the *TgOI* (Fig. 3B). None of the F_1_s from the 25_17A and 25_17B founder families offspring exhibited the appearance of single-copy fish, whereas only a few F_1_ embryos obtained from 25_17C founder family seemed to be single-copy (Fig. 3B, S3).

Thus, application of the competitive dilution strategy for Tol2 transgenesis significantly increased the proportion of single-insertion transgenics in F_1_ generation. Notably, the total copy number of *TgOI* and *BFP* transgene in offspring from the Mix_17 injection founder families was comparable to the copy number determined in the offspring of F_0_ fish injected with 25 pg of pMK17 plasmid (Fig. 3C,D).

### The ubiquitin promoter is subject to position effects

Our *TgOI* cassettes are under the control of the *ubb* promoter, known for constitutive transgene expression throughout all developmental stages and adult organs (Mosimann et al., 2011). Yet, while analysing the single-insertion offspring from the different founder families, we observed clear differences in mRFP expression (Fig. 4). Apart from expected differences in overall strength of expression, we also noted variation in brightness of fluorescence in different tissues, including a strong myotome-specific enhancer-trap (Fig. 4) and a lateral line enhancer-trap (line not saved). Thus, even ubiquitous promoters can be subject to strong position effects, further underscoring the importance of using single-insertion transgenic lines.

**Fig. 4.**
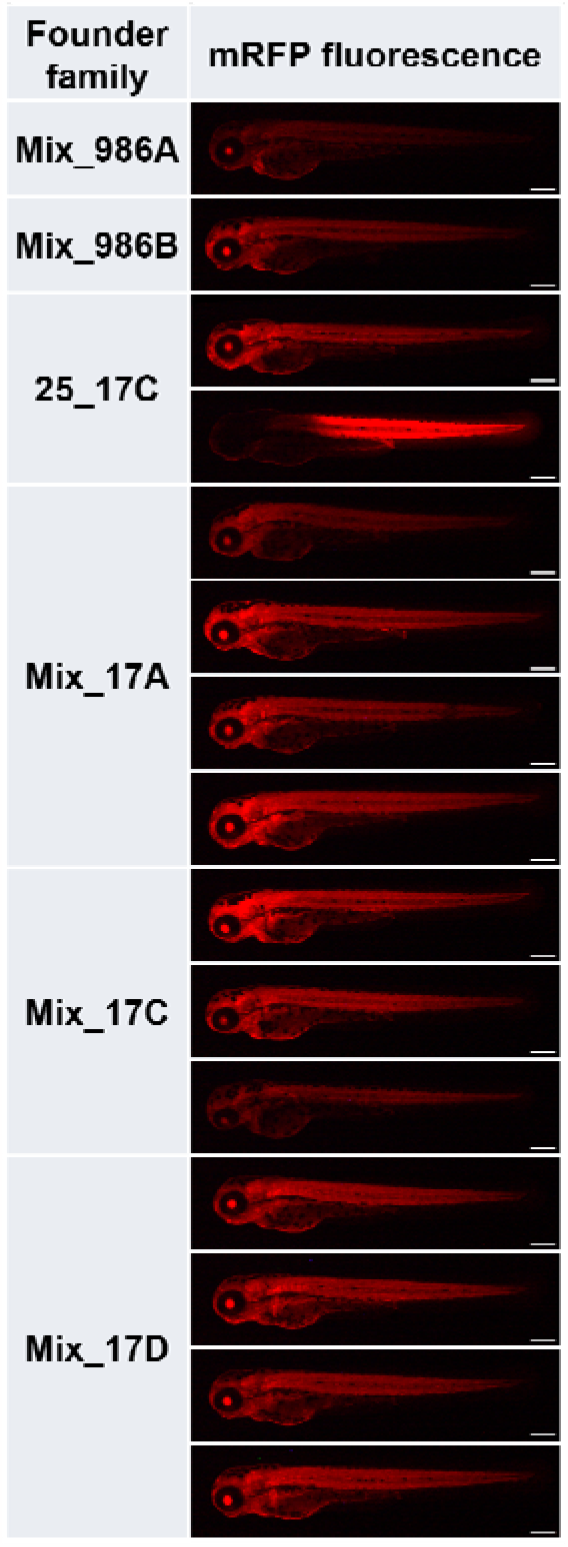
The widely used ubiquitin (ubb) promoter is subject to position effects. Single-insertion transgenic lines display differences in mRFP expression pattern. All images underwent consistent enhancement to facilitate clearer visualization of expression differences. Scale bars: 250μm.

Our competitive dilution strategy significantly enhances the standard Tol2-mediated transgenesis protocol producing a high percentage of single-insertion transgenic zebrafish already in the F_1_ generation. While we used a lens-BFP marked transgene as the competitor, it can be replaced with any other robust fluorescently marked Tol2 transgene. Our modified protocol reduces animal usage and saves time while also enhances reproducibility by facilitating publication and distribution of single-insertion transgenic lines.

## MATERIALS AND METHODS

### Zebrafish maintenance

Zebrafish (*Danio rerio*) were used in accordance with Temple University Institutional Animal Care and Use Committee (IACUC) guidelines under the approval from protocol numbers ACUP 4354, ACUP 4709 and/or in accordance with an approved license of Animal research ethics committee (Lithuania) No. B5-10954 and permission for the experimentation from the State Food and Veterinary Service of Lithuania, No. G2-231. Male and female breeders from 3-12 months of age were used to generate fish for all experiments. Strains generated in this study are: *Tg(ubb:loxP-mRFP-STOP-loxP-nup35-eGFP-BLRP)*^*tpl133*^, *Tg(ubb:loxP-mRFP-STOP-loxP-nup35-eGFP-BLRP)*^*tpl134*^, *Tg(ubb:loxP-mRFP-STOP-loxP-tcf21-3xHA)*^*vln15*^. Embryos and larvae were anesthetized with 4 mg/ml MS-222 (Sigma-Aldrich, E10521) dissolved in ddH_2_O and diluted in egg water for handling when necessary.

### Generation of plasmids

Details on plasmid construction are available upon request.

### Tol2 mRNA synthesis

pT3TS-Tol2 (Balciunas et al., 2006) was linearized with XbaI, purified using GeneJET Gel Extraction Kit (Thermo Scientific, K0692) and transcribed using the T3 mMESSAGE mMACHINE™ *in vitro* transcription kit (Invitrogen, AM1348). Transcribed mRNA was purified using either GeneJET RNA Purification Kit (Thermo Scientific, K0732) or RNeasy MinElute Cleanup Kit (Qiagen, 74204), diluted to 40 ng/μl in RNase-free water (Invitrogen, 10977035) and divided into 2 μl aliquots. Aliquots were stored at − 80 °C.

### Microinjections

The *TgOI* plasmids (pDB986 or pMK17) were mixed with diluting plasmid (pDB815) in a 1:4 mass ratio to achieve a final DNA concentration of 10 ng/μl. An 8 μl of plasmids mix was combined with 2 μl aliquot of Tol2 mRNA and approximately 3 nl of mix (25 pg of plasmids/Tol2 mRNA per embryo) were injected into the yolks of single-cell-stage zebrafish embryos as described previously (Balciuniene and Balciunas, 2013). For pMK17 injection, 8 μl of 10 ng/μl plasmid was combined with 2 μl aliquot of Tol2 mRNA and approximately 3 nl of mix (25 pg of plasmid/Tol2 mRNA per embryo) were injected into the yolks of single-cell-stage zebrafish embryos as described previously (Balciuniene and Balciunas, 2013).

### Screening for germline transmission

Injected embryos were visually screened at 3 dpf for mRFP fluorescence using Leica DM5500 B or Zeiss AxioImager microscopes. mRFP-positive embryos were raised to adulthood. Adult F_0_ fish were incrossed. Offspring were visually screened for mRFP and BFP expression using Leica DM5500 B or Zeiss AxioImager microscopes.

### Imaging and image processing

All imaging was performed on a Leica DM5500 B microscope using the same parameters. Live imaging was conducted with a HC PL FLUOTAR 2,5 ×/0,07 objective (Leica) mounting 3 dpf embryos in 3% methylcellulose (Thermo Scientific, 258111000). After imaging, each individual embryo was placed in a separate well of a 48-well plate and allowed to develop until 5 dpf. Imaging data were processed identically in ImageJ/FIJI (National Institutes of Health).

### Transgene copy number determination

5 dpf embryos were euthanised in 4 mg/ml MS-222 (Sigma-Aldrich, E10521) solution, transferred to 1,5 ml microcentrifuge tube and frozen. Genomic DNA (gDNA) was isolated by digesting sample in DNA extraction buffer (0,1 M Tris pH-9 (Fisher BioReagents, BP152), 0,1 M NaCl (Fisher BioReagents, BP358), 0,05 M EDTA (Sigma-Aldrich, ED2SS), 0,2 M Sucrose (Fisher Chemical, S/8560/65), 0,5% SDS (Carl Roth, CN30.3)) with 0,2 mg/ml Proteinase K (Thermo Scientific, EO0491). gDNA was precipitated using 8 M Potassium Acetate (Sigma-Aldrich, P-1147) and isopropanol (Sigma-Aldrich, 24137-M) followed by washing with 70% ethanol (Sigma-Aldrich, 32221-M). After that gDNA was incubated with RNase A (Thermo Scientific, EN0531) to remove any traces of RNA, purified, and dissolved in TE buffer. To precisely determine *TgOI* copy number we performed qPCR using primers specific for *mRFP*, with *tpl133Tg* heterozygotes serving as a single copy control. For copy number determination of the diluting transgene we used primers specific for *BFP*, using heterozygous *tb ×5a*^*tpl58R*^ gene trap line, containing the *gamma-crystalline:BFP* expression cassette (Grajevskaja et al., 2018), as a single copy control. *thyroglobulin precursor* (TG) gene was used as a reference locus since it is present in the zebrafish genome in a stable number (Wang et al., 2007; Zhang et al., 2019). qPCR was performed on Rotor-Gene Q (QIAGEN) using Maxima SYBR Green qPCR Master Mix (Thermo Scientific, K0253) without ROX passive dye. qPCR cycling conditions were as follows: 2 min at 50 °C, 10 min at 95 °C, followed by 40 cycles of 15 s at 95 °C, 30 s at 60 °C, and 30 s at 72 °C. Relative copy number of the transgenes was determined using the 2^−ΔΔCt^ method (Livak and Schmittgen, 2001). Primers sequences for qPCR are summarized here: mRFP-Fw: TGAACGGCCACGAGTTCGAGAT; mRFP-Rv: TCATCACGCGCTCCCACTTGAA; BFP-Fw: ACGTAAACGGCCACAAGTTC; BFP-Rv: GGGGGTGTTCTGCTGGTAG; TG-Fw: GGTATCTACGGTTTAGAATCCGATACATG; TG-Rv: TGGAGTCCTCTTTACTAGTCTCAGTC.

## Acknowledgements

We thank Dr. Frank L. Conlon (University of North Carolina) for sharing the *nup35-eGFP-BLRP* cassette.

