## Supplementary Figures 1-3 for "Rapid generation of single-insertion transgenics by Tol2 transposition in zebrafish"

| Founder family | Fish line | mRFP fluorescence |
|---|---|---|
| Mix_986A | *tpl133Tg* | 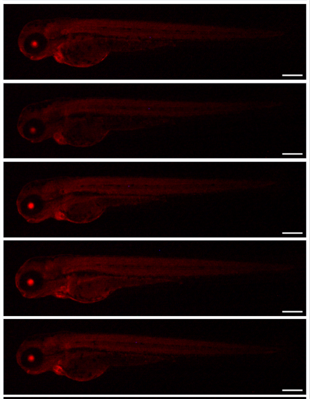 |
| Mix_986B | *tpl134Tg* | 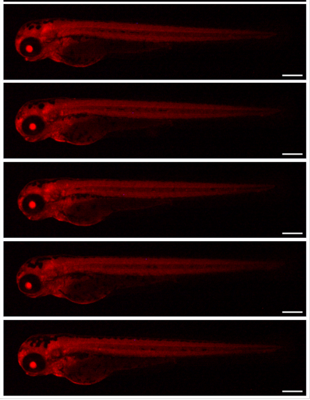 |

**Fig. S1.** Transgene expression in *tpl133Tg* and *tpl134Tg* embryos. Scale bars: 250µm.

| Founder family | *TgOI* copy no. | *BFP* copy no. | mRFP fluorescence | BFP fluorescence |
|---|---|---|---|---|
| Mix_17A | 1,4 | 2,4 | | |
| | 2,8 | 3,1 | | |
| | 1,3 | 2,8 | | |
| | 1,3 | 2,1 | | |
| | 1,0 | 0,0 | | |
| | ND | ND | | |
| | ND | ND | | |
| | ND | ND | | |
| | ND | ND | | |
| | ND | ND | | |
| | ND | ND | | |

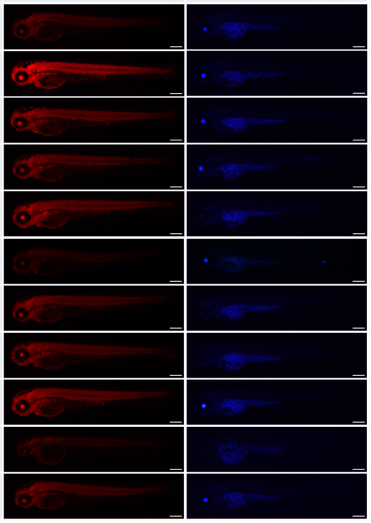

| Founder family | *TgOI* copy no. | *BFP* copy no. | mRFP fluorescence | BFP fluorescence |
|---|---|---|---|---|
| Mix_17C | 2,4 | 3,2 | | |
| | 1,1 | 2,2 | | |
| | 1,1 | 3,0 | | |
| | 2,7 | 4,2 | | |
| | 1,1 | 4,5 | | |
| | ND | ND | | |
| | ND | ND | | |
| | ND | ND | | |
| | ND | ND | | |
| | ND | ND | | |
| | ND | ND | | |
| | ND | ND | | |
| | ND | ND | | |

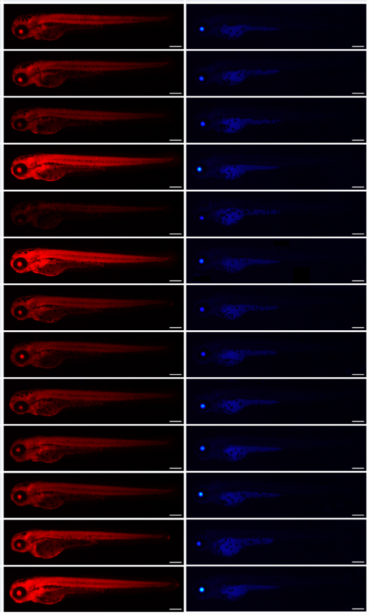

| Founder family | *TgOI* copy no. | *BFP* copy no. | mRFP fluorescence | BFP fluorescence |
|---|---|---|---|---|
| Mix_17B | 2,0 | 7,9 | | |
| | 5,2 | 11,8 | | |
| | 2,7 | 12,5 | | |
| | 3,9 | 10,4 | | |
| | 8,3 | 6,9 | | |
| | ND | ND | | |
| | ND | ND | | |
| | ND | ND | | |
| | ND | ND | | |
| | ND | ND | | |
| | ND | ND | | |
| | ND | ND | | |
| | ND | ND | | |
| | ND | ND | | |
| | ND | ND | | |
| | ND | ND | | |
| | ND | ND | | |
| | ND | ND | | |
| | ND | ND | | |
| | ND | ND | | |

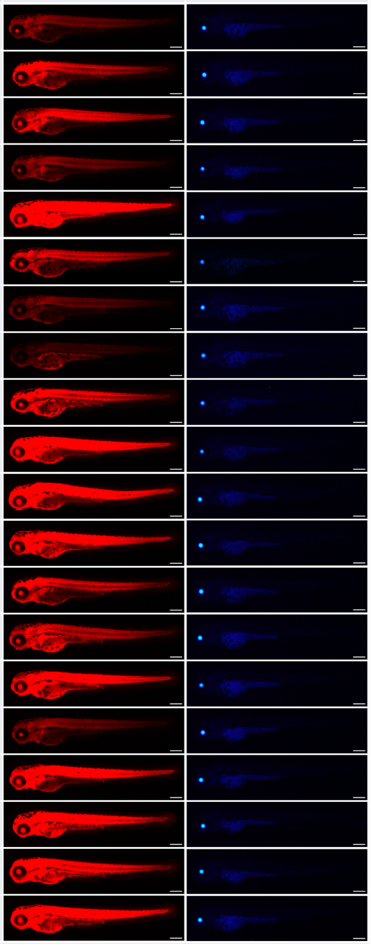

| Founder family | *TgOI* copy no. | *BFP* copy no. | mRFP fluorescence | BFP fluorescence |
|---|---|---|---|---|
| Mix_17D | 1,3 | 0,0 | | |
| | 1,2 | 1,0 | | |
| | 1,5 | 0,8 | | |
| | 1,1 | 0,0 | | |
| | 1,2 | 1,1 | | |
| | ND | ND | | |
| | ND | ND | | |
| | ND | ND | | |
| | ND | ND | | |
| | ND | ND | | |
| | ND | ND | | |

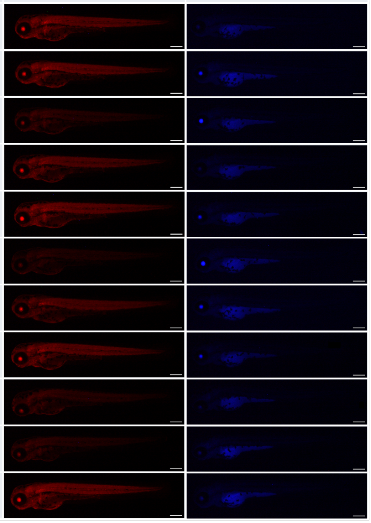

**Fig. S2.** Estimated *TgOI* and *BFP* transgene copy number and mRFP/BFP expression in $F_1$ embryos from 4 different founder families injected with a mix of pMK17/pDB815 plasmids. Scale bars: 250µm.

| Founder family | *TgOI* copy no. | mRFP fluorescence |
|---|---|---|
| 25_17A | 20,3 | |
| | 9,4 | |
| | 14,7 | |
| | 22,9 | |
| | 16,3 | |
| | ND | |
| | ND | |
| | ND | |
| | ND | |
| | ND | |
| | ND | |
| | ND | |
| | ND | |
| | ND | |
| | ND | |

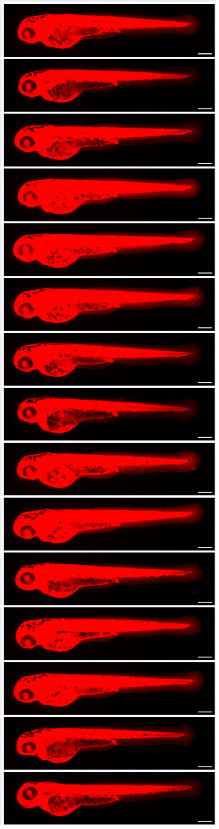

| Founder family | *TgOI* copy no. | mRFP fluorescence |
|---|---|---|
| 25_17B | 6,5 | |
| | 6,7 | |
| | 5,3 | |
| | 2,7 | |
| | 1,7 | |
| | ND | |
| | ND | |
| | ND | |
| | ND | |
| | ND | |
| | ND | |
| | ND | |
| | ND | |
| | ND | |
| | ND | |

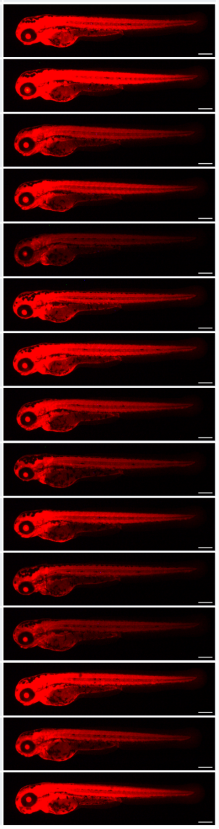

| Founder family | *TgOI* copy no. | mRFP fluorescence |
|---|---|---|
| 25_17C | 11,5 | |
| | 7,5 | |
| | 3,9 | |
| | 1,2 | |
| | 1,2 | |
| | ND | |
| | ND | |
| | ND | |
| | ND | |
| | ND | |
| | ND | |
| | ND | |
| | ND | |
| | ND | |
| | ND | |
| | ND | |

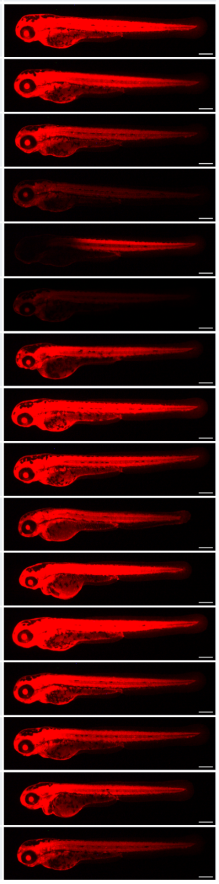

**Fig. S3.** Estimated *TgOI* copy number and mRFP expression in $F_1$ embryos from 3 different founder families injected with 25 pg of pMK17 plasmid. Scale bars: 250µm.
